# QTLseqr: An R package for bulk segregant analysis with next-generation sequencing

**DOI:** 10.1101/208140

**Authors:** Ben N. Mansfeld, Rebecca Grumet

## Abstract

1

Next Generation Sequencing Bulk Segregant Analysis (NGS-BSA) is efficient in detecting quantitative trait loci (QTL). Despite the popularity of NGS-BSA and the R statistical platform, no R packages are currently available for NGS-BSA. We present QTLseqr, an R package for NGS-BSA that identifies QTL using two statistical approaches: QTL-seq and G’. These approaches use a simulation method and a tricube smoothed G statistic, respectively, to identify and assess statistical significance of QTL. QTLseqr, can import and filter SNP data, calculate SNP distributions, relative allele frequencies, G’ values, and log_10_(p-values), enabling identification and plotting of QTL. The source code is available at https://github.com/bmansfeld/QTLseqr.

**Core ideas:**

- An R package that performs Next Generation Sequencing Bulk Segregant Analysis was developed
- Two methods for analysis are provided: QTL-seq and G’
- The QTLseqr package is quick and produces publication quality figures and tables

## 2 Introduction

Since the early 1990’s, Bulk Segregant Analysis (BSA) has been a valuable tool for rapidly identifying markers in a genomic region associated with a trait of interest (Giovannoni et al., 1991; Michelmore et al., 1991). BSA is amenable to any type of codominant markers, including single nucleotide polymorphism (SNP) markers. This has allowed for the adaptation of this technology for use with next-generation sequencing (NGS) reads. The recent reduction in cost of NGS has further contributed to the increased use and development of this and similar methods [thoroughly reviewed by Schneeberger, (2014)].

The NGS-BSA procedure is performed by establishing and phenotyping a segregating population and selecting individuals with high and low values for the trait of interest. DNA from these individuals is pooled into high and low bulks which are subject to sequencing and single nucleotide polymorphism (SNP) calling, thus mitigating a need to develop markers in advance. In bulks selected from F_2_ populations, SNPs detected in reads derived from regions not linked to the trait of interest should be present in ~50% of the reads. However, SNPs in reads aligning to genomic regions closely linked to the trait should be over‐ or under-represented depending on the bulk. Thus, comparing relative allele depths, or SNP-indices (defined as the number of reads containing a SNP divided by the total sequencing depth at that SNP) between the bulks can allow quantitative trait loci (QTL) identification (Takagi et al., 2013).

In plant breeding research, the main pipeline used for BSA, termed QTL-seq, was developed by Takagi et al. (2013) and has been widely used in several crops for many traits (e.g. Das et al., 2014; Lu et al., 2014; Win et al., 2016; and many others). Takagi and colleagues define the Δ(SNP-index) for each SNP as the difference of the low value bulk SNP-index from the high value bulk SNP-index. They suggest averaging and plotting Δ(SNP-indices) over a sliding window. Regions with a Δ(SNP-index) that pass a confidence interval threshold, as calculated by a statistical simulation, should contain QTL. The algorithm described by Takagi et al. was released as a pipeline written in a combination of bash, pearl and R, meant to perform all tasks from trimming, processing and cleaning raw reads to plotting Δ(*SNP-index*) plots.

An alternate analytical pipeline to evaluate statistical significance of QTL from NGS-BSA was proposed by Magwene et al. (2011). A modified G statistic is calculated for each SNP based on the observed and expected allele depths and smoothing this value using a Nadaraya-Watson, or tricube smoothing kernel (Nadaraya, 1964; Watson, 1964). This smoothing method weights neighboring SNPs’ G statistic by their relative distance from the focal SNP such that closer SNPs receive higher weights. Using the smoothed G statistic, or G’, Magwene et al. allow for noise reduction while also addressing linkage disequilibrium (LD) between SNPs. One advantage to this method is that p-values can be estimated for each SNP using non-parametric estimation of the null distribution of G’. This provides a clear and easy-to-interpret result as well as the option for multiple testing corrections.

Due to its general ease-of-use, multi-system compatibility, open-source nature, and ease of package distribution, the statistical programming language R (https://www.r-project.org/) has rapidly established its status as the tool-of-choice for computational biology analyses (Tippmann, 2014). As no scripts were released to facilitate G’ analysis, and no R packages were available for performing NGS-BSA, we developed the QTLseqr package with the goal of making both QTL-seq and G’ methods accessible to plant breeders and geneticists. QTLseqr can be easily installed and is highly configurable, allowing the user control of many parameters and the type of analysis performed. QTLseqr rapidly performs genome-wide calculations and simulations required for either method, and produces publication ready plots and tables, allowing for easy identification of putative QTL regions. The full source code is available at https://github.com/bmansfeld/QTLseqr.

## 3 Features and methods

### 3.1 Overview

A straight forward pipeline for analysis was designed with the plant breeder and geneticist in mind: 1) Import SNP data, 2) Filter SNPs that may complicate analysis, 3) perform bulk segregant analyses, 4) plot results and 5) export the data. A vignette with a step-by-step guide is available at https://github.com/bmansfeld/QTLseqr/vingettes.

### 3.2 Data import and filtering

QTLseqr imports SNP data, from GATK’s *VariantsToTable* function (Van der Auwera et al., 2013), as a data frame where each row is a SNP and each column is a descriptive field. For each SNP, the total reference allele frequency, per bulk SNP-index, and Δ(SNP-index) are calculated. To help reduce noise and improve results, the *filterSNPs()* function offers options for filtering SNPs based on reference allele frequency, total read depth, per bulk read depth and genotype quality score. Filtering by read depth can help eliminate SNPs with low confidence due to low coverage, or SNPs that may be in repetitive regions and thus have inflated read depth. The initial number of SNPs, number of SNPs filtered per step, total number of SNPs filtered, and remaining number are reported.

### 3.3 Bulk segregant analyses

Both methods, QTL-seq or G’ methods, are comparable in their ability to detect QTL, but differ in sensitivity based on their different defined thresholds. The methods are somewhat complimentary and calculating Δ(SNP-index) is informative in both analyses, as the contributing parent of the QTL may be inferred by the Δ(SNP-index) value in the region. QTLseqr can perform NGS-BSA using either or both methods and results may be compared to confirm identified QTL.

#### 3.3.1 The QTL-seq approach

QTL-seq analysis is performed using the *runQTLseqAnalysis()* function, which first counts the number of SNPs within the set window bandwidth. The subsequent analysis is derived from the original pipeline of Takagi et al. (2013) with some minor changes. 1) Instead of using a uniform or “rectangular” window as originally suggested, we opt for a tricube-smoothed Δ(SNP-index) calculated similarly to G’, which smooths-out noise, while accounting for LD between SNPs (Supplemental equations 1-4). 2) To fully take advantage of R’s rapid vectorized calculations, scripts have been rewritten to perform the simulations that define read-depth-based confidence intervals at each SNP position (Supplemental Fig. S1). Several simulation parameters are user-configurable including: the population type (F_2_ or RIL), simulated read depth, number of bootstrapped replications, and a filter threshold for simulated reads. The user can then extract QTL, defined as contiguous genomic regions whose absolute tricube-smoothed Δ(SNP-index) values are higher than the simulated intervals, using the *getSigRegions()* and *getQTLToble()* functions, described below.

#### 3.3.2 The G’ approach

For the G’ approach, the primary analysis steps are performed by *runGprimeAnalysis()* which initially calculates the G statistic (Supplemental equations 5-9) for each SNP. It then counts the number of SNPs within the set window bandwidth and estimates the tricube-smoothed G’ and Δ(SNP-index) values of each SNP within that window (Supplemental equations 4, 10).

One benefit of the G’ method is that p-values and genome-wide Benjamini-Hochberg (Benjamini and Hochberg, 1995) false discovery rate (FDR), adjusted p-values are calculated for each SNP. As it is close to being log-normally distributed, p-values can be estimated from the null distribution of G’, which assumes no QTL. To this end, G’ values from QTL regions are temporarily removed from the full set, so that mean and variance of the null distribution of G’ may be estimated. Magwene *et al.* (2011) suggest using Hampel’s rule (an outlier filtering approach, [Davies and Gather, 1993]) to filter out these regions. However, with the data we tested (Yang et al., 2013) this method failed to filter any values (Supplemental Fig. S2). Alternatively, filtering G’ values in regions of high absolute Δ(SNP-index) is a data driven method effective in identifying and filtering potential QTL. We find that this approach is successful for estimating p-values and offer it, alongside Hampel’s rule, as an option for p-value calculation.

### 3.4 Plotting and exporting result

QTLseqr has two main plotting functions for quality control and data visualization. The *plotGprimeDist()* function can be used to plot the G’ distribution as a check to assess the validity of the analysis (Supplemental Fig. S2). The *plotQTLStats()* function is used for plotting the number of SNPs/window, the tricube-weighted Δ(*SNP-index*) and *G’* values, or the ‐log_10_(p-value) (Fig. 1).

**Figure 1.**
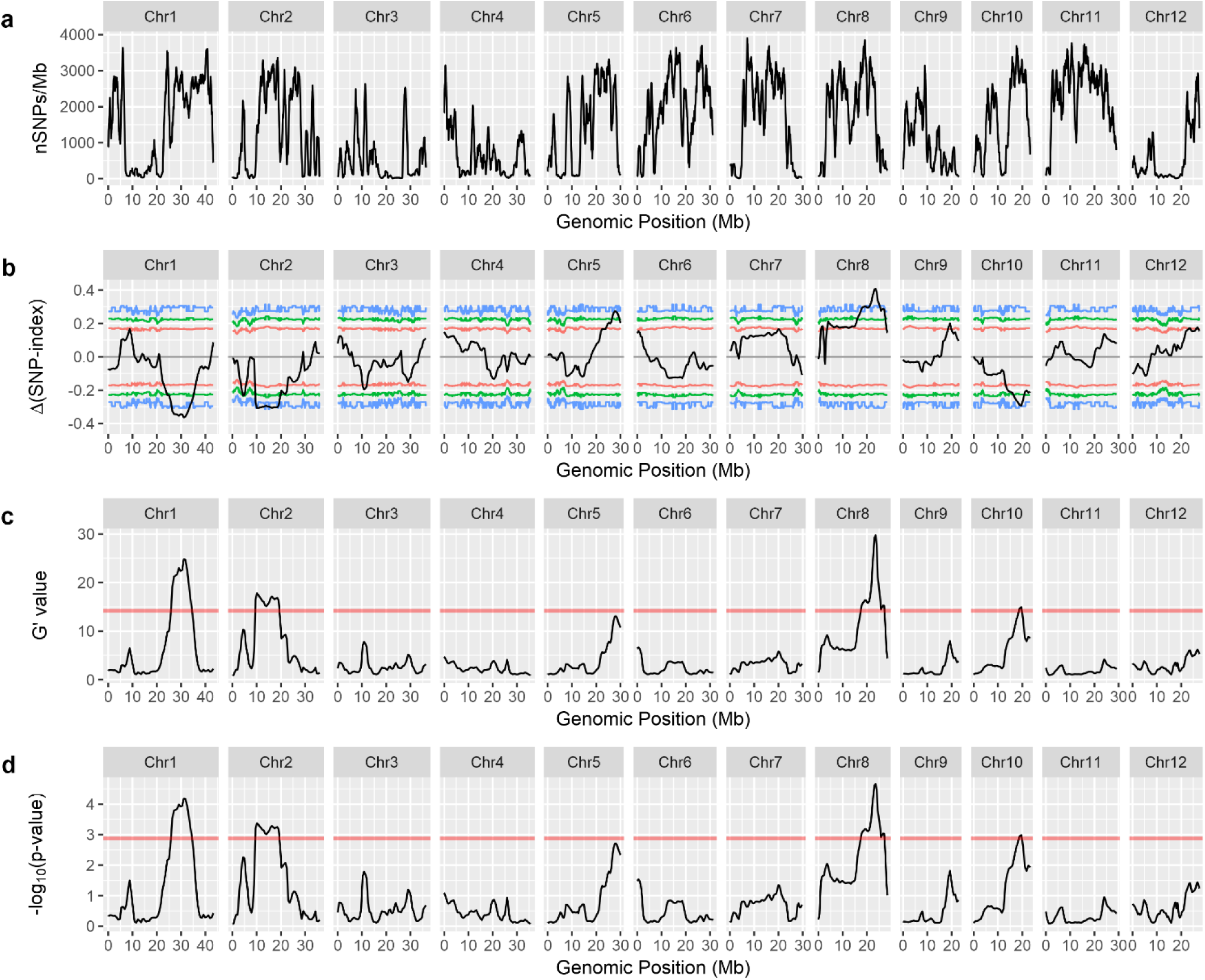
Quantitative trait loci for rice seedling cold tolerance identified by QTLseqr. Plots produced by the *plotQTLStats()* function with a 1 Mb sliding window: Distribution of SNPs in each smoothing window (a). The tricube-smoothed Δ(SNP-index) and corresponding two-sided confidence intervals: 95% (red), 99% (green), and 99.9% (blue) (b). The tricube-smoothed G’ value (c). Another, more familiar way to display QTL, is using the ‐log_10_(p-value) which is derived from the *G’* value, (d). In (c) and (d) the genome-wide false discovery rate of 0.01 indicated by the red line.

QTLseqr functions are available for extracting, summarizing and reporting of significant QTL regions. The *getSigRegions()* function will produce a list in which each element represents a QTL region. The elements are subsets of the original data frame supplied. Any contiguous region with a q-value above the set alpha (G’ method), or absolute Δ(SNP-index) above the requested confidence interval (QTL-seq method) will be returned. If there is a dip below the threshold the region will be split to two elements. The getQTLTable will summarize those results in a table and can output a comma-separated value file, if requested (e.g. Table 1).

**Table 1.**
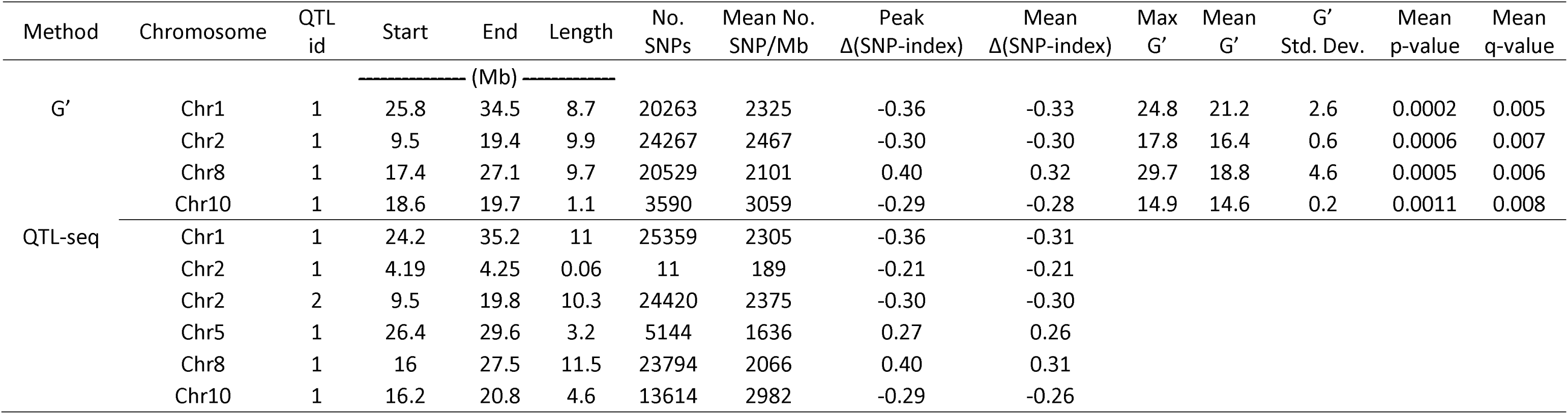
Quantitative trait loci (QTL) identified by QTLseqr in test data (Yang et al., 2013). QTL were defined as regions with a q-value above the false discovery rate of 0.01 or a Δ(SNP-index) above a confidence interval of 99% for G’ or QTL-seq, respectively.

## 4 Implementation and results

As a test of the validity and efficacy of our package functions, and to compare the two analysis methods, we tested QTLseqr’s ability to reproduce results described by Yang et al. (2013), a BSA study which utilized the G’ approach to identify loci for seedling cold tolerance in rice. Raw reads were downloaded from the NCBI Short Read Archive, aligned to the v7 Nipponbare genome (http://rice.plantbiology.msu.edu/) and SNPs were called as described in the GATK “Best Practices” (https://software.broadinstitute.org/gatk/best-practices/). Detailed methods are available in Supplemental Material.

QTLseqr was successful at reproducing the analysis performed by Yang and colleagues, confirming QTL on chromosomes 1, 2, 8, and 10 using either analysis method. Figure 1 shows the putative QTL identified, as output by the *plotQTLStats()* function. The results of our analyses are summarized in Table 1 as provided by the *exportQTLTable()* function in QTLseqr. While both methods were successful in identifying the same regions as QTL, the boundaries of each region largely depended on the confidence interval or FDR rate that was chosen. Using a confidence interval of 99% with the QTL-seq method was not as stringent as using a FDR of 0.01 in the G’ method. As such, the QTL-seq method detected a second narrow region on Chromosome 2, as well as a region on Chromosome 5, which was also originally reported by Yang et al. (2013).

## 5 Conclusion

The QTLseqr package provides a fast and straightforward tool for plant breeders and other scientists to perform NGS-BSA using either QTL-seq or G’ analysis methods. Data from the identified QTL can be exported for downstream analysis and summarized in publication ready figures and tables.

## 7 Acknowledgments and funding

We thank the members of the Michigan State University, Horticulture Department’s Joint Lab group for critical reading of the manuscript as well as Drs. Robin Buell and Robert VanBuren for other feed back and advice. This work was in part supported by the National Institute of Food and Agriculture, US Department of Agriculture, under award number 2015-51181-24285 and MSU Project GREEEN.

